# Induction of cytoplasmic dsDNA and cGAS-STING immune signaling after exposure of breast cancer cells to X-rays or high energetic carbon ions

**DOI:** 10.1101/2024.07.23.604756

**Authors:** C. Totis, N. B. Averbeck, B. Jakob, M. Schork, G. Volpi, D.F. Hintze, M. Durante, C. Fournier, A. Helm

**Author notes:** TRON - Translational Oncology at the University Medical Center of the Johannes Gutenberg University gGmbH, Mainz, Germany. Radiobiology Unit, Research and Development Department, CNAO National Center for Oncological Hadrontherapy, Pavia, Italy. Paul-Ehrlich-Institut, Division of Allergology, Section Research Allergology, Langen, Germany (D.F. Hintze). **Corresponding author:** Alexander Helm. Author responsible for statistical analysis: Cristina Totis.

## Abstract

Radiotherapy can trigger activation of the cGAS-STING axis via cytoplasmic dsDNA fragment induction. The activation of cGAS–STING initiates innate immune signaling mediated by interferon type-I that can contribute to eradicate the malignancy. The effect was shown to depend on the fractionation scheme employed. We hypothesized that the innate immune response can also depend on radiation quality because densely ionizing radiation, such as carbon ions, have different effects on DNA lesion quality. We show here that carbon ions induced a significantly higher yield of cytosolic dsDNA fragments per unit dose as compared to photons in an *in vitro* 4T1 breast cancer model. The higher efficiency also translated in expression and release of interferon-β by the tumor cells. Cytoplasmic dsDNA fragments as well as interferon-β release increased with doses up to 24 Gy and no differences for a fractionation scheme (3x8 Gy) were found as compared to the single high doses of photons. In conclusion, we found that the release of interferon-β after radiation is increasing with the radiation dose up to 20 Gy and that carbon ions have the potential to elicit a strong innate immune signaling.

## Introduction

Triple-negative breast cancer remains a challenge for treatment due to its resistance to standard treatments, especially chemotherapy and radiotherapy (1). As for radiotherapy, the location of the tumor represents a further challenge as lung and heart are sensitive organs at risk and may receive a certain dose during treatment, increasing the risk for late adverse cardiovascular effects (2). Due to a higher precision and better sparing of healthy tissue, particle therapy using protons is considered an alternative and has been shown to be safe and feasible, reducing the dose to non-target structures while optimizing target coverage, but long-term clinical data are yet scarce (3). Carbon ion radiotherapy (CIRT) is also attractive because it offers comparable precision as compared to protons plus an increased biological effectiveness as compared to photons (4). This increased biological effectiveness results in an improved control of radioresistant tumors and may be also more immunogenic (5). Immunogenicity of the local radiotherapy treatment is important for combination with immunotherapy, such as immune checkpoint inhibitors (6). Vanpouille-Box and colleagues (7) demonstrated radiation-induced activation of the cGAS/STING pathway with a subsequent type-I-interferon response, initially triggered by the occurrence of dsDNA fragments in the cytoplasm. Additionally, the authors found a discontinuous and non-linear dose response, where the balance between presence of dsDNA fragments and expression of *Trex1*, which clears the cytoplasm from the fragments, determines the effectiveness. The most effective dose range in that context was 8 to 12 Gy, and a fractionated 3x8 Gy regimen was superior to a single high dose of ≥20 Gy based on the same mechanistic interplay (7). In this study, we investigate the potential of CIRT in a murine triple-negative breast cancer model in terms of overcoming radioresistance and especially in triggering immunogenicity via the occurrence of cytoplasmic dsDNA fragments and a subsequent type-I-interferon response.

## Material and Methods

### Cell lines and reagents

The mammary carcinoma cell line 4T1 was purchased from ATCC® (CRL-2539 ™) and cultured in Roswell Park Memorial Institute medium (RPMI 1640 + GlutaMAX^TM^-I, Gibco) supplemented with 10% fetal bovine serum (Sigma-Aldrich®) and 1% Penicillin Streptomycin (10 000 Units/mL Penicillin, 10 mg/mL Streptomycin, Gibco). The TS/A mammary adenocarcinoma cell line was purchased from EMD Millipore (SCC177) and cultured in Dulbecco’s Modified Eagle’s Medium (DMEM+ GlutaMAX^TM^-I, Gibco) with high glucose containing 10% fetal bovine serum (Sigma-Aldrich®) and 1% Penicillin Streptomycin (10 000 Units/mL Penicillin, 10 mg/mL Streptomycin, Gibco). All cell lines were maintained in culture with 5% CO_2_ at 37° C and routinely screened for *Mycoplasma* (PCR Mycoplasma test kit, ITW Reagents). For some of the experiments, the cGAS inhibitor (RU.521, InvivoGen) was used at 20 µM, the STING inhibitor (H-151, InvivoGen) at 1.5 µM. Both were added immediately before irradiation.

### Irradiation

Cells were seeded in 25 cm^2^ tissue culture flasks and at 70-80% of confluency were exposed to X-rays. For quantifying cytoplasmic dsDNA, 7.5 x 10^5^ 4T1 cells per 3.5 cm Petri dishes were seeded 24 h before irradiation. For X-ray irradiation, we used the vertical irradiation cabinet (X-ray generator Isovolt DSI (Seifert, Ahrensberg), 7 mm beryllium, 1 mm aluminum, and 1 mm copper filter system (250 kV/16 mA) at a dose rate of 2 Gy/min at the GSI Helmholtz Center for Heavy Ion Research (Darmstadt, Germany). Carbon-ion irradiation (^12^C^6+^) was performed at the Marburg Ion Beam Therapy center (MIT, Marburg, Germany, MIT-2022-03) or at the SIS18 at the GSI Helmholtz Center for Heavy Ion Research, irradiating 5x10^6^ cells per dose (220 MeV/u; dose-averaged linear energy transfer (LET) of 75 keV/μm, 4 cm SOBP). Immediately after irradiation the cells were seeded in 25 cm^2^ tissue culture flasks. For both radiation types physical doses from 2 to 24 Gy were delivered. Additionally, a fractionation regimen of 8 Gy given in 3 consecutive days (3x8 Gy) was performed for X-rays.

### Clonogenic cell survival

Clonogenic survival of 4T1 cells was assessed using the colony forming assay as described in (8). After irradiation, cells were counted and plated in triplicates into 25 cm^2^ tissue culture flasks. The appropriate number of cells was seeded aiming at the statistically significant formation of 100 colonies. The cells were fixed and stained with methylene blue solution after one week of incubation. A colony was counted if composed of at least 50 cells.

### Immunofluorescence, confocal microscopy, and quantification of cytosolic dsDNA

For immunofluorescence staining, cells were fixed in 2 % formaldehyde and permeabilized with 0.1 % Triton X-100 for 10 min and blocked with 0.4 % BSA in PBS for 20 min. All antibodies were diluted in 1x PBS, 0.4 % BSA: αdsDNA (host: mouse; Abcam, ab27156) was used at a concentration of 1 µg/ml and αmouse Alexa Fluor 488 (host: goat; Invitrogen, A-11017) at a concentration of 5 µg/ml. In order to visualize the cytoplasmic area, we stained F-actin with Phalloidin-iFluor 647 Conjugate (Cayman, 20555). DNA was counterstained with 4′,6-diamidino-2-phenylindole (DAPI) (1µg/ml, AppliChem, A1001). Upon mounting in Slow Fade Diamond Antifade (Invitrogen, S36963) the cells were imaged at a Nikon Eclipse Ti microscope. The exposure time for the immunostained dsDNA signal was optimized for the cytosolic signal, which in most cases caused overexposure of the nuclear dsDNA signal. To quantify the cytosolic dsDNA signals within the images, we used the in-house developed software ImageD (versions v1_7_0 and v2_0_4, plugin “cytoplasmic dsDNA detection” https://github.com/DavidEilenstein/ImageD). With this software, dsDNA signals ≥ 5 px were quantified as cytoplasmic dsDNA foci, if they were located outside the DAPI stained area (nucleus) and inside the phalloidin stained area (cytoplasm). dsDNA foci counted in 0 Gy samples (background) were subtracted from dsDNA foci numbers at doses > 0 Gy. For X-rays, three to six independent experiments were performed (exception: 2 Gy; one experiment). For carbon-ions, two independent experiments were performed (exception: 20 Gy; one experiment). Per experiment, at least 53 cells (20 Gy X-rays) and up to 1425 cells (0 Gy carbon ions) were analyzed per dose.

### Gene expression

Total RNA from cell pellet was extracted with RNeasy Mini Kit (Qiagen) according to the manufacturer’s instructions, including a DNase digestion step (RNase-free DNase set, Qiagen). 2 µg of extracted RNA were used to perform cDNA synthesis, according to the manufacturer’s instructions (RevertAid First Strand cDNA Synthesis Kit, Thermo Fisher Scientific). Subsequently, the cDNA was loaded in a quantitative polymerase chain reaction (qPCR) plate (Sarstedt) with Fast SYBR^TM^ Green Master Mix (Thermo Fisher Scientific) and bioinformatically validated specific primers (Gapdh, QT00199388; Rpl13a, QT00267197; Ifnb1, QT00249662; Trex1, QT00288967; QuantiTect Primer Assay, Qiagen). The final volume was 20 µL and the cycling conditions were a holding stage of 95°C for 15 min, followed by 40 cycles of denaturation at 95°C for 15s, annealing at 60°C for 30s, extension at 72°C for 20s. A melt curve analysis was performed to ensure the specificity of the amplicons. The qPCRs were run using QuantStudio^TM^ 3 system (Thermo Fisher Scientific) and the results collected with QuantStudio^TM^ Design & Analysis Software (version 1.5.2, Thermo Fisher Scientific). The fold change of the genes of interested was calculated with the 2^(-ΔΔCt)^ method, normalizing the Ct values for 2 reference genes (*Gapdh, Rpl13a*).

### ELISAs

IFN-β release was quantified with ELISA kit (PBL Assay Science, PBL42410-1) according to manufacturer’s instructions. An additional dilution of the standard curve was necessary due to the low concentration of the analyte in the controls. For cGAMP assessment, cells were pre-treated with M-PER® Mammalian Protein Extraction Reagent (Thermo Fisher Scientific) following manufacturer’s instructions, according to the sample preparation recommendations of the ELISA kit (2’3’ cGAMP ELISA kit, Cayman Chemical, 501700). The ELISA was performed according to manufacturer’s specifications. Absorbance (450 nm) was measured with BioTek EL808 Microplate Reader and Gen5 (version1.11) software.

### Bulk RNA Sequencing

Sequencing was performed by Arraystar Inc. (Rockville, MD, USA). In brief, 1-2 µg of total RNA of 4T1 and TS/A cells were used to prepare the sequencing library. The libraries were sequenced on Illumina NovaSeq 6000 instrument. Fold change (cutoff 1.5), p-value (≤ 0.05), and FPKM (≥ 0.5 mean in one group) were used for filtering differentially expressed genes and transcripts. The Z-score was calculated for some Genes Of Interest (GOI). For further details, please see the supplementary section.

### Statistics

Unless otherwise specified, the experiments were performed in 3 biological replicates for X-rays and carbon ions, and data are presented as mean ± standard deviation (SD). The data were analyzed with GraphPad Prism software (version 9.3.1) and statistically significant differences between X-rays and carbon ions were determined using unpaired two-tailed *t*-test or two-way analysis of variance (ANOVA) with Tukey’s multiple comparisons test and considered significant for *P*-values ≤ 0.05, unless indicated elsewise. For further information on the performed fits, we refer to the supplementary section.

## Results

### Carbon ions are more effective than X-rays in killing triple negative breast-cancer cells and inducing cytoplasmic dsDNA fragments

Carbon ions (C-ions) are more effective than X-rays in 4T1 cell killing and the relative biological effectiveness (RBE) is about 2 (Fig. 1). We then measured the induction of dsDNA in the cytoplasm after radiation. Carbon ions resulted in significantly more cytoplasmic dsDNA foci per cell when compared to photons 24h following exposure (Fig. 2A and B). We also investigated the dose response of *Trex1* expression, which is known to remove dsDNA fragments from the cytoplasm. Expression of *Trex1* increased continuously with dose at 24h from X-ray exposure, and slightly higher values were observed after carbon ions at high doses compared to the same X-ray doses (Fig. 2C). Expression of *Trex1* remained similar both for amplitude and dose response at 48h (Fig. 2D). To corroborate such dose response, we investigated the *Trex1* expression in TS/A cells, confirming the increase with dose (Suppl. Fig. 1A and B). Consistent with the steady increase for dsDNA foci over dose, *Trex1* seemed not to have a threshold dose from which on its expression would strongly increase to remove fragments. In order to link these observations with the activity of cGAS, we measured the concentration of cGAMP (a product of cGAS activation) at 24h following exposure to photons or carbon ions (Suppl. Fig. 2). It slightly increased with dose with carbon ions being in tendency (not significantly) more efficient.

**Figure 1.**
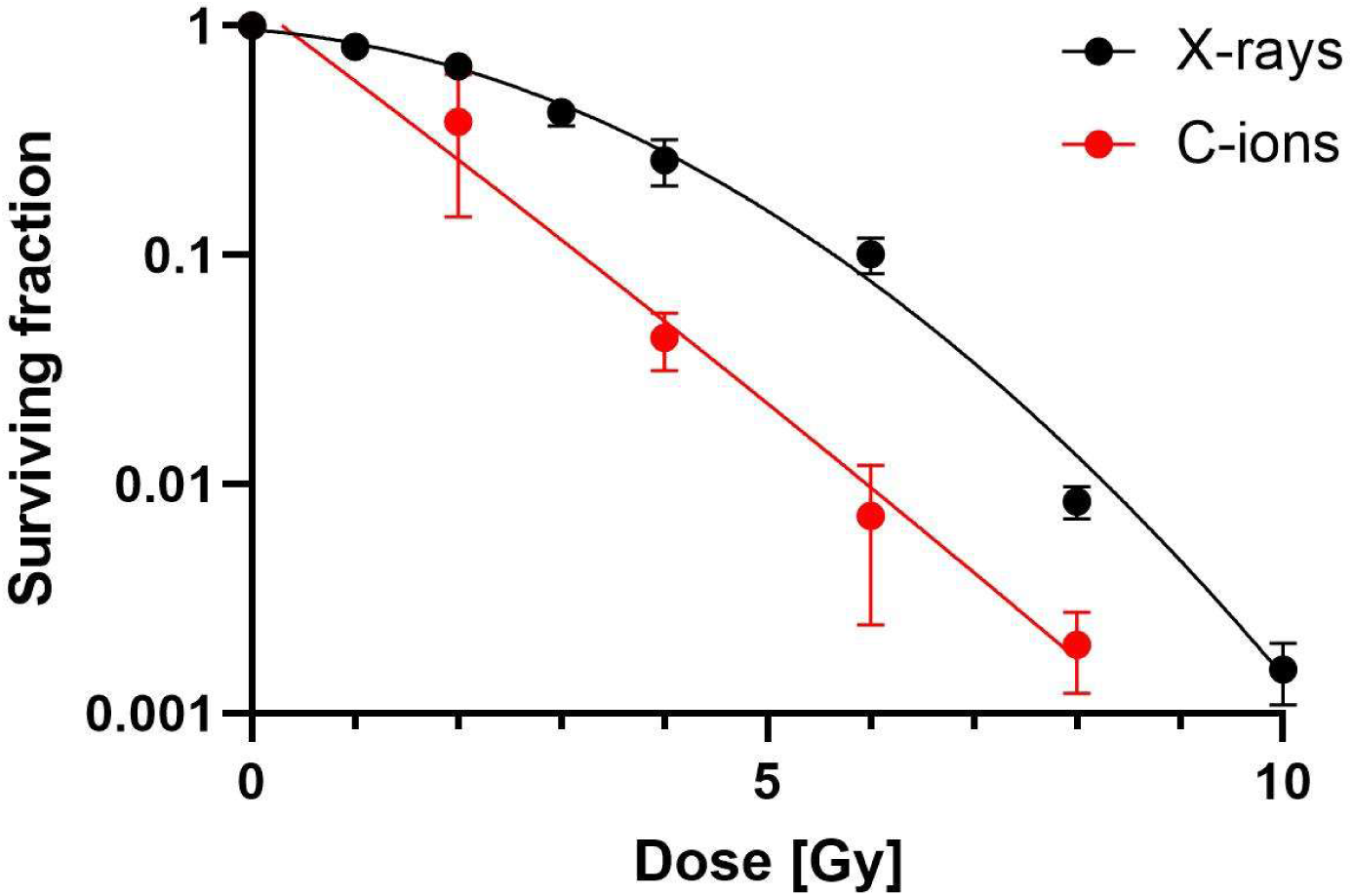
Clonogenic cell survival. Colonies of surviving 4T1 cells were counted following exposure to X-rays or carbon ions and plotted as surviving fraction. Curves were fitted using the linear quadratic model. The relative biological efficiency (RBE) was calculated to be roughly 2 at a level of 10% survival. Data for X-rays were derived from Reppingen et al. (23).

**Figure 2.**
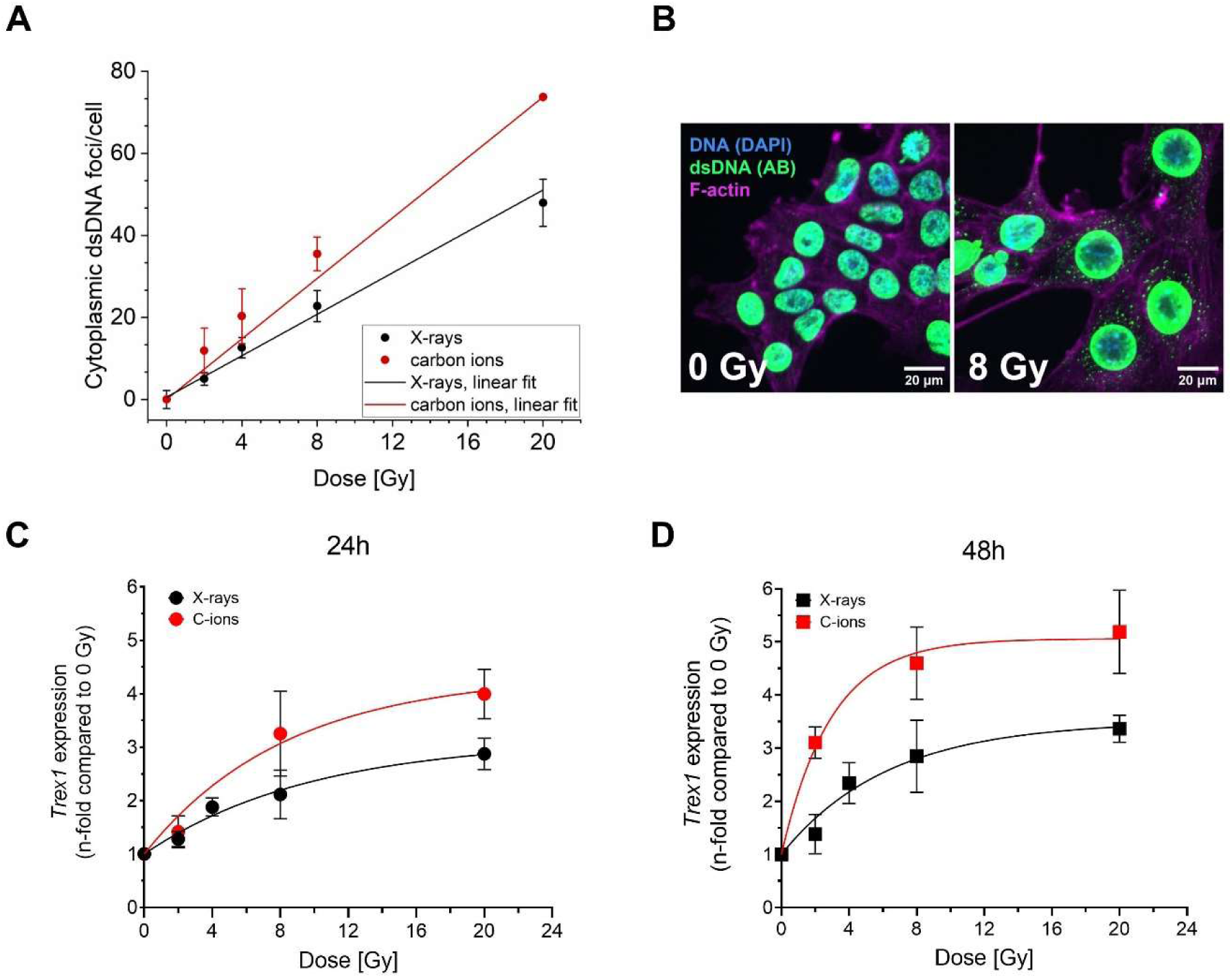
Foci of cytoplasmic dsDNA fragments and expression of *Trex1*. (A) Foci of dsDNA fragments per cell in dependence of the dose. Numbers of cytoplasmic dsDNA foci of non-irradiated cells were subtracted. 4T1 cells were irradiated with X-rays or carbon ions. Carbon ions resulted in significantly higher amounts of foci 24 h after irradiation (see supplement for details on the statistical analysis). (B) Representative images of 4T1 cells 24 h after 0 Gy or 8 Gy X-ray irradiation. dsDNA was immunofluorescence stained (AB), F-actin as a cytosol marker was stained with phalloidin, and DNA was also counterstained with DAPI. dsDNA signals outside the DAPI stained nucleus but inside the phalloidin stained area were counted as cytoplasmic dsDNA foci. Expression of *Trex1* in 4T1 cells increases with dose at 24 h (C) and 48 h (D). The curves tend to reach a plateau at high doses. Comparison of the trends was performed using an exponential function (see Supplements). The difference between X-rays and C-ions is not statistically significant at 24 h but is statistically significant at 48 h (see Supplementary materials).

### Carbon-ion irradiation results in an elevated interferon-β release indicating its potential for increased immunogenicity

We performed bulk RNA sequencing 24h after exposure to X-rays or carbon ions in order to study changes in gene expression based on the different radiation qualities (Fig. 3). Generally, carbon ions led to a higher number of genes differentially expressed than X-rays (Fig. 3A, and Suppl. Fig. 3A and B) when compared to their respective controls. Figure 3B displays a set of selected genes related to immunogenic signaling. Generally, the expression of these genes increases following 8 or 20 Gy of X-rays or carbon ions, with a slightly higher expression for 20 Gy carbon ions in some of the genes. This was found both for genes related to immunoactivation and immunosuppression and confirmed in TS/A cells upon exposure to X-rays when compared to 4T1 cells (Suppl. Fig.4A and B). As for damage-associated molecular patterns (DAMPs), no coherent pattern of dose response was found. More in detail, focusing on genes related to the type-I-interferon response, *Ifnb1* expression was below the limit of quantification with this method at 24h and is therefore not displayed here. *Trex1* expression pattern over dose confirmed the dose response found by qRT-PCR (see Fig. 2). Similar dose responses were also found for *Cxcl10*, a gene coding for a cytokine (CXCL10) released upon auto- and paracrine type-I-interferon signaling. Interestingly, *Cxcr3*, a gene for a receptor of CXCL10 appeared not to be increased following carbon-ion exposure.

**Figure 3.**
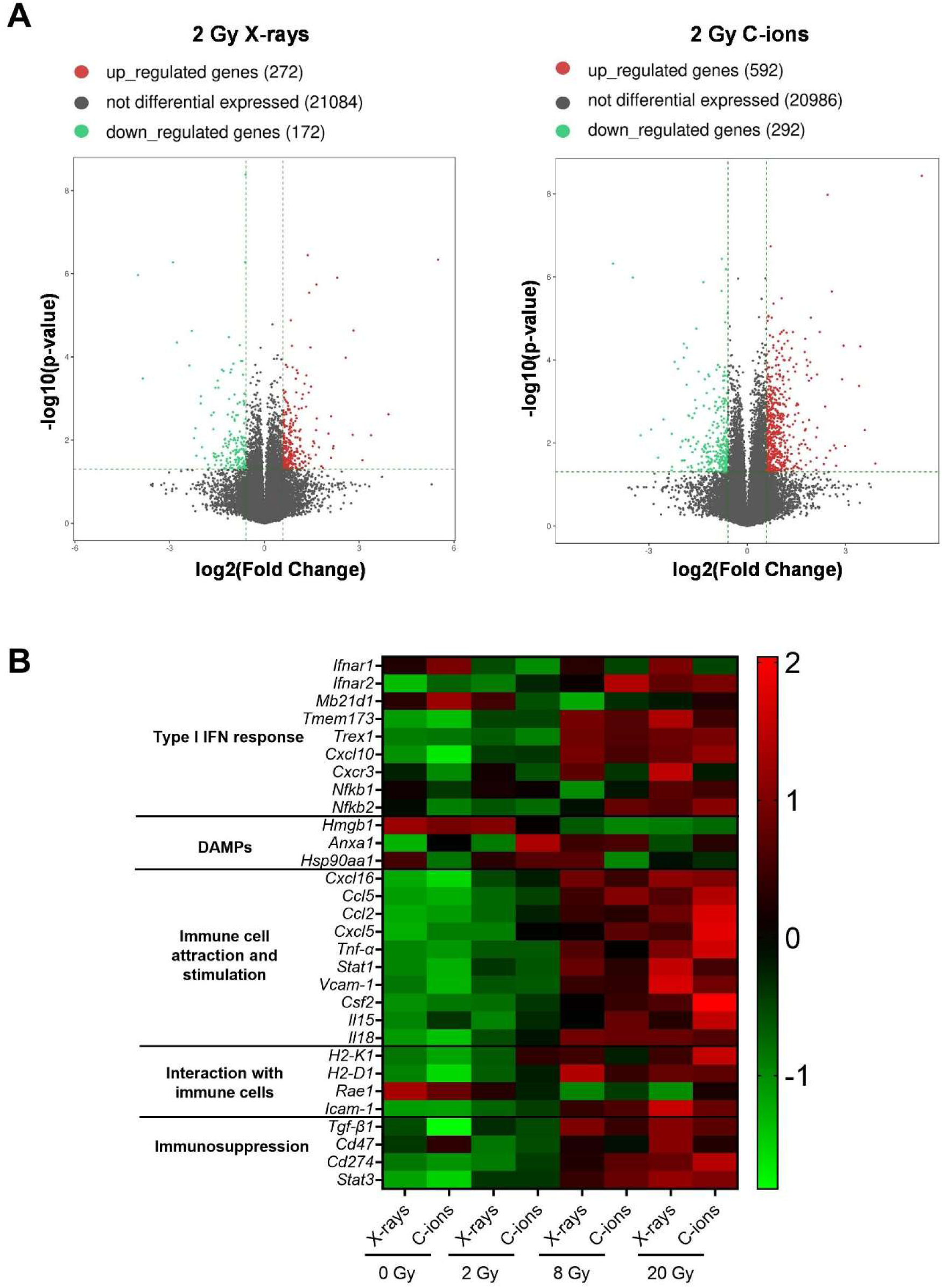
Effects of X-rays or carbon ion irradiation on differential expression of genes in 4T1 cells. Volcano plot of differentially expressed genes at 24h after 2 Gy irradiation with X-rays (left) or carbon ions (right) compared to the respective non-irradiated samples (0 Gy). Significantly upregulated genes and downregulated genes are depicted in red and green, respectively, and non-significant genes are shown in gray (A). Z-score hierarchical clustering heat map visualization of the expression of some genes of interest (rows) across different doses and radiation qualities (columns). Colors represent scaled expression levels, with green for low expression and red for high expression (B).

As IFN-β is a crucial factor in the type-I-interferon response, yet was not detectable in bulk RNA sequencing, we measured the expression of *Ifnb1* with qRT-PCR and the release of IFN-β at 24h following exposure. *Ifnb1* gene expression showed an increase with dose and carbon ions being more effective than photons (Fig. 4A), which is in line with the dose response of cytoplasmic dsDNA fragments (Fig. 2A). The release of IFN-β at 24h was generally low with little differences between the two radiation types (Fig. 4B). Therefore, we investigated both expression and release also 48h following exposure (Fig. 4C and 4D). The levels of both were elevated with respect to 24h and the dose response stayed similar. Comparable dose responses following X-ray exposure were again confirmed in TS/A cells (Suppl. Fig. 1C to F).

**Figure 4.**
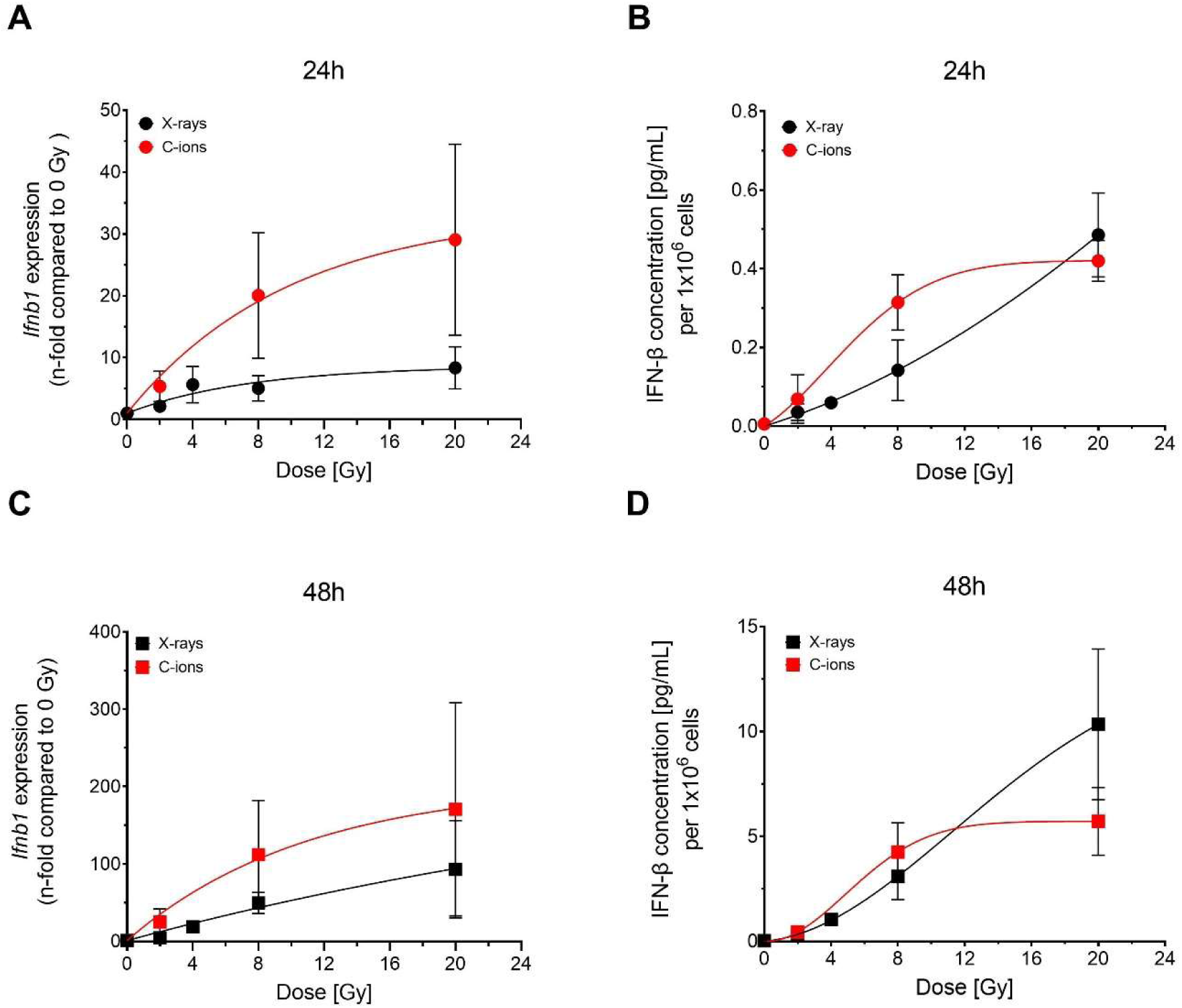
Expression of *Ifnb1* and release of IFN-β. Expression of *Ifnb1* was measured at 24h (A) and 48h (C) after exposure to X-rays or carbon ions. At similar time points the release of IFN-β (B and D) was assessed. With the exception of 20 Gy in release, where a plateau was reached, carbon ions were more efficient in tendency, but not significantly (see Supplementary material) in 4T1 cells.

To demonstrate the link of the effects to the activation of the cGAS/STING pathway, we next investigated its downstream signal, i.e. the expression of the *Ifnb1* gene following the inhibition of each cGAS or STING. While exposure to X-rays resulted in an increased expression of *Ifnb1* at 24h, application of cGAS or STING inhibitors diminished (cGAS inhibitor, Suppl. Fig. 5A) or eradicated (STING inhibitor, Suppl. Fig. 5B) the effect, demonstrating the involvement of the cGAS/STING pathway.

Based on the observation that the expression of *Ifnb1* and the release of its gene product is higher 48h than 24h after irradiation, we speculated that a differential response in IFN-β release might occur only at later time points. Therefore, we measured the amount of released IFN-β in dedicated samples daily on a time course of up to 144h (Fig.5A and 5B). Indeed, at 72h following irradiation, we detected strong differences (p=0.0004 and p=0.0019 for 8 Gy and 24 Gy, respectively) in the release of IFN-β between photons and carbon ions. The release was not only stronger but also more persistent following exposure to carbon ions (Fig. 5B), indicating an increased potential for a robust immunogenic signal.

**Figure 5.**
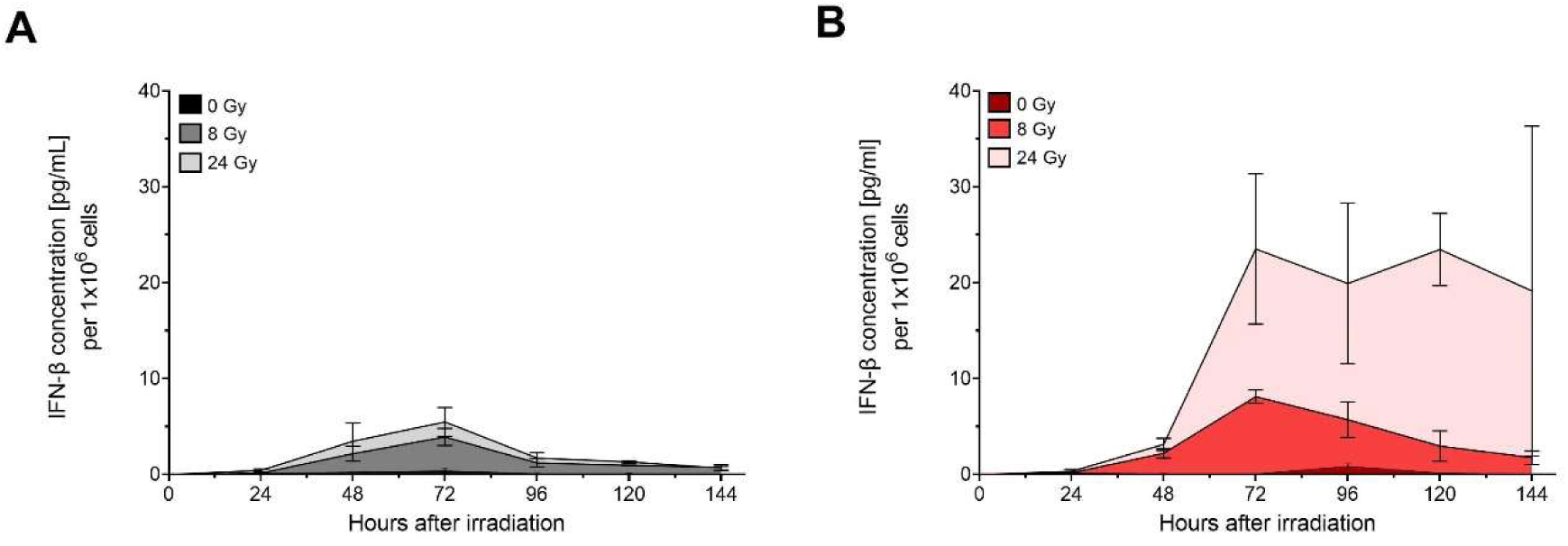
Release of IFN-β over time. The release of IFN-β was measured up to 144h after exposure to 8 Gy or 24 Gy X-rays (A) or carbon ions (B) in 4T1 cells. At 72h post irradiation, carbon ions cause a significantly higher release of IFN-β than X-rays (8 Gy p=0.0004, 24 Gy p=0.0019, unpaired two-tailed t-test, N=5 X-rays, N=3 carbon ions).

### A single high dose is comparatively effective to a fractionated scheme following exposure to photons

We investigated whether a hypofractionated regimen of 3x8 Gy was superior to a single high dose of photons (24 Gy) (Fig. 6A-F). Our results do not support the hypothesis that fractionation increases cGAS-STING pathway activation. While we indeed found a reduced level of *Trex1* expression at 24h and 48h after exposure to 3x8 Gy (Fig. 6A), this did not translate in higher amounts of cytoplasmic dsDNA foci per cell (Fig. 6B). While slightly higher amounts of *Ifnb1* expression and IFN-β release were measured at 24h after exposure to 3x8 Gy X-rays, the effect did not persist as at 48h after exposure the effect was found inversed (Fig. 6C and D). This is confirmed by IFN-β release over time, in which following 3x8 Gy there is a lower release than after a single dose of 24 Gy (Fig. 6E). Measuring cGAMP as a surrogate of cGAS activation, again, we found a similar effect (Fig. 6F). Measuring *Trex1* and *Ifnb1* expression as well as IFN-β release in TS/A cells confirmed little to no differences between 24 Gy and 3x8 Gy (Suppl. Fig. 6 A-C).

**Figure 6.**
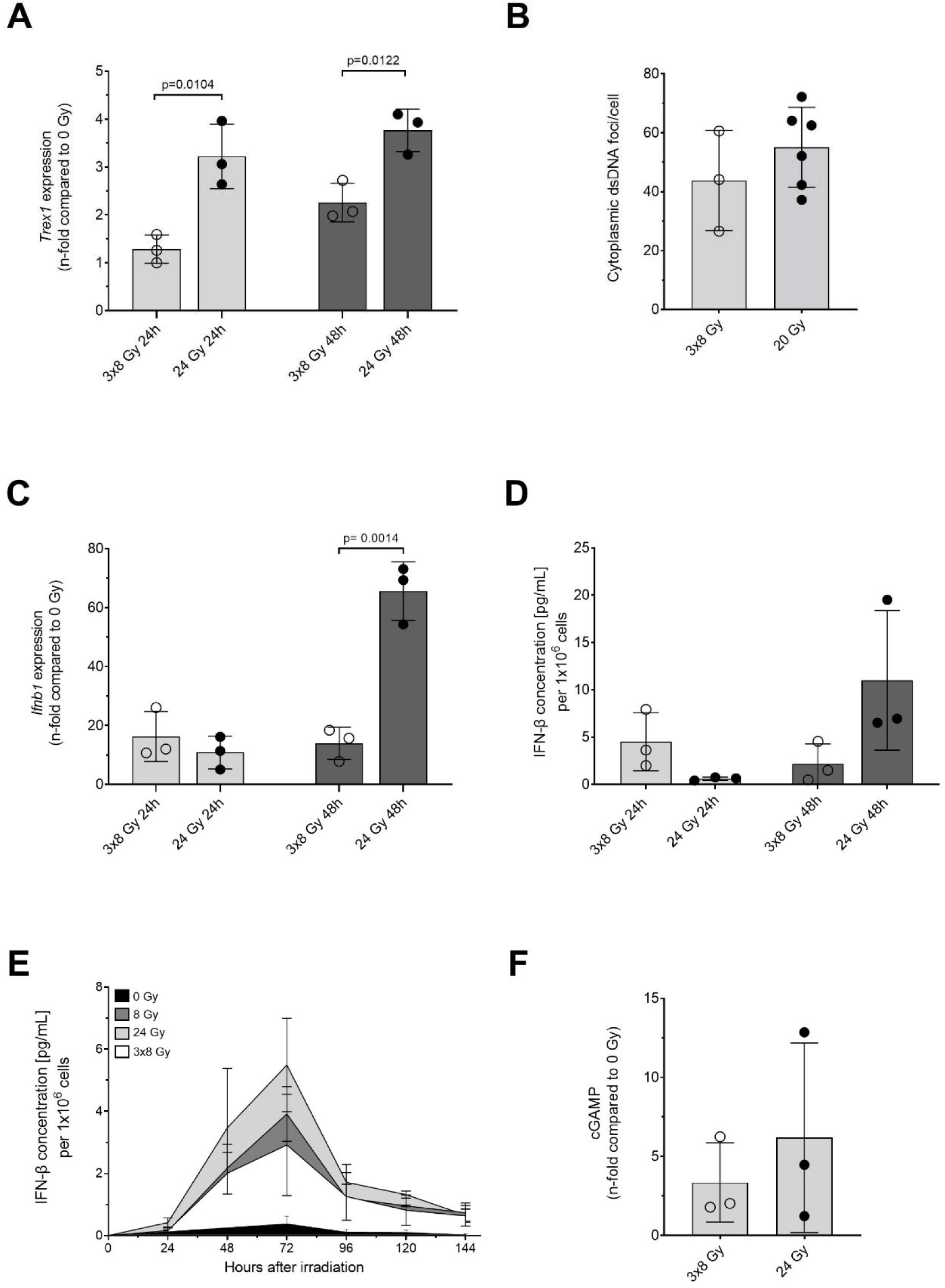
Comparison between single high doses and a 3x8 Gy hypofractionation scheme in 4T1 cells. The endpoints tested for the dose response curves were also performed comparing a single high dose of 20 or 24 Gy to a 3x8 Gy hypofractionation scheme at 24h or 48h after exposure to X-rays. More in detail, *Trex1* expression (A), cytoplasmic dsDNA foci 24 h after X-rays (B), *Ifnb1* expression (C), IFN-β release (D), IFN-β release over time (E) and cGAMP 24h after X-rays (F) were measured. Significances were tested using an unpaired two-tailed t-test. (E) Data for single doses are derived from Fig. 5 A.

## Discussion

Carbon ion radiotherapy (CIRT) is increasingly used worldwide as it features improved precision in beam delivery and an increased biological effectiveness, which allows treatment of radioresistant tumors with close vicinity to sensitive organs (9). Additionally, CIRT is considered to feature an increased immunogenicity, which renders it a good match for immunotherapies and the treatment of metastatic cancer disease (10–12). Clinical trials are currently being set up to prove this hypothesis (13, 14). Using the poorly immunogenic murine breast cancer model 4T1 (15), we investigated the immunogenicity of carbon ions with respect to the type-I-interferon response triggered by the cGAS/STING axis, which in turn is activated by the presence of dsDNA foci in the cytoplasm (16).

We hypothesized that carbon ions can be more effective with respect to the occurrence of cytoplasmic dsDNA fragments due to their peculiar features of DNA damage induction (17). Indeed, our data indicated that carbon ions resulted in an increased number of cytoplasmic dsDNA foci, which translated in an increased release of IFN-β, pointing to a higher potential for a strong type-I-interferon response. Furthermore, the increased efficiency of carbon ions with respect to clonogenic cell survival, hence overcoming radioresistance of a triple-negative breast cancer model, provides additional rationale for the application of CIRT in breast cancer therapy (4). In this context, the dual role of interferons has to be considered as a sustained or prolonged IFN signaling may result in adverse effects related to therapy resistance (18, 19). Of note, our data indicated lower expression of the gene coding for CXCR3 for carbon ions as compared to X-rays. Coexpression of CXCL10/CXCR3 on tumors was reported to be adverse with respect to metastasis development and to be a marker for poor clinical outcome. Autocrine signaling of the Cxcl10/Cxcr3 axis was shown to promote tumor cell growth, motility and metastasis (20). Interestingly, a reduced metastatic potential for CIRT has already been reported in other pre-clinical models (11, 21).

We failed to reproduce the previously reported discontinuous dose response, favoring a single dose in the range of 8 to 12 Gy over a single high dose due to the mechanistic interplay between the occurrence of cytoplasmic dsDNA fragments and *Trex1* expression (7). In both murine breast cancer models, 4T1 and TS/A cells, used as well by Vanpouille-Box and colleagues, our data show an increase of the signal with dose throughout several endpoints. The former also reported best responses for a 3x8 Gy regimen as compared to single doses, again based on the above-mentioned mechanistic dualism. Our study instead resulted in comparable signals for a single dose of 24 Gy vs. a 3x8 Gy regimen. Despite using the same cell lines, different results between laboratories can be explained with the application of different materials such as fetal bovine serum or different assays. Moreover, the method of choice for quantification of cytoplasmic dsDNA in our study is based on the assessment of foci per cell, rather than recovering a total signal in a group of cells, which could partly account for the differences.

Nonetheless, we found a coherent increase with dose in all endpoints. Vanpouille-Box and colleagues showed that their results translated *in vivo* when the respective doses were applied in a mouse model in combination with immune checkpoint inhibitors, corroborating their hypothesis of superiority of a fractionated regimen over a single high dose (7). Our study lacks an *in vivo* translation as of now, which could verify this dose response and the superiority of CIRT over photons. Our data also indicate the importance of assessing several time points and later than the commonly used 24h after exposure in *in vitro* studies, as a differential release of cytokines occurred only at later time points, likely related to the radiation-induced cell cycle block.

If verified in further pre-clinical models, our results would argue for the application of higher doses than the classical 2 Gy fractionation scheme, as a dose of 8 Gy consistently resulted in enhanced signals. Also, despite not being superior to high doses of ≥20 Gy, we found a 3x8 Gy scheme at least comparable to 8 Gy. In fact, hypofractionated schemes applying 3x8 Gy or schemes in that range are increasingly used in clinical settings (22). Since carbon ions mostly performed better than X-rays at 8 Gy, our data encourage the application of 8 Gy or hypofractionated regimens also for CIRT, and indeed clinical studies with such regimens are already set up (13).

Finally, since the occurrence of dsDNA fragments in the cytoplasm is the base of the type-I-interferon response, and since carbon ions resulted in a higher amount of dsDNA foci, the question of by which mechanism the fragments travel to the cytoplasm and how can that be differentially affected through carbon ions (17) becomes crucial but remains to be addressed. In summary, our data support the application of CIRT in the context of immunotherapy combinations but underpin that further research is required at the same time to better exploit its potential advantages.

## Data sharing statement

All data generated and analyzed during this study are included in this published article (and its supplementary files).

## Acknowledgements

We thank M. Scholz and T. Friedrich for planning and realizing the particle exposure of cells at GSI. We thank S. Lerchl and M.C. Martire for helpful discussions as well as D. Kraft and A. Benzer for skilful assistance.

