## Supplementary Material for "Induction of cytoplasmic dsDNA and cGAS-STING immune signaling after exposure of breast cancer cells to X-rays or high energetic carbon ions"

### Supplement

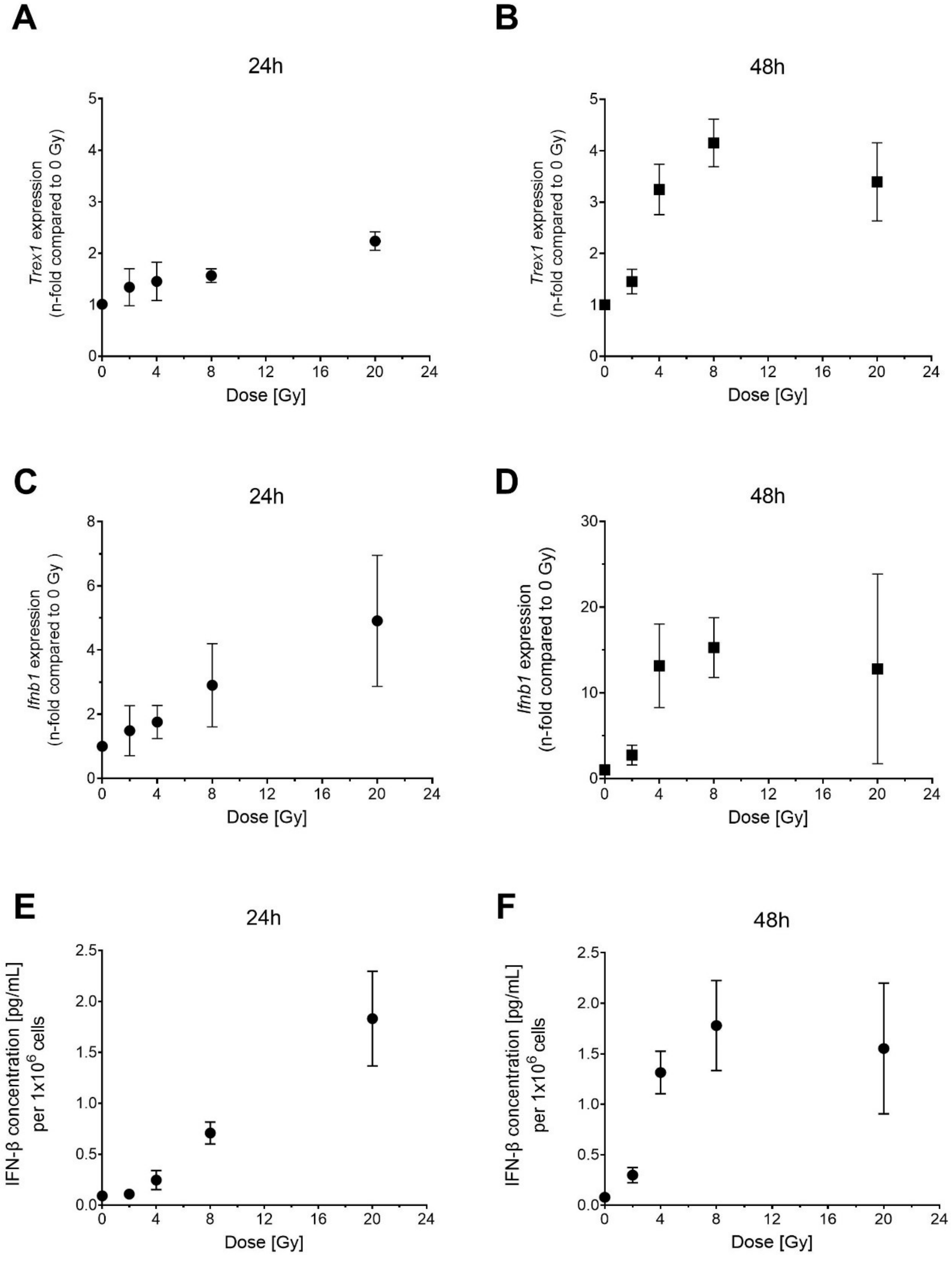

**Supplementary Figure 1**. **Expression of *Trex1, Ifnb1* and release of IFN-β in TS/A cells.** Expression of *Trex1* (A 24h, B 48h) and *Ifnb1* (C 24h, D 48h) as well as release of IFN-β (E 24h, F 48h) were measured following exposure to X-rays.

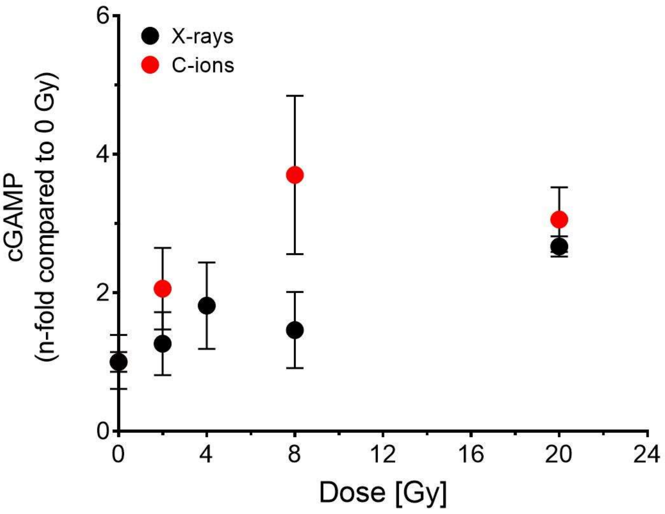

**Supplementary Figure 2**. **Cellular concentration of cGAMP as a surrogate for cGAS activity in 4T1 cells.** The concentrations were measured at 24h after exposure to X-rays or carbon ions. No significant differences were found between radiation qualities (unpaired two-tailed t-test).

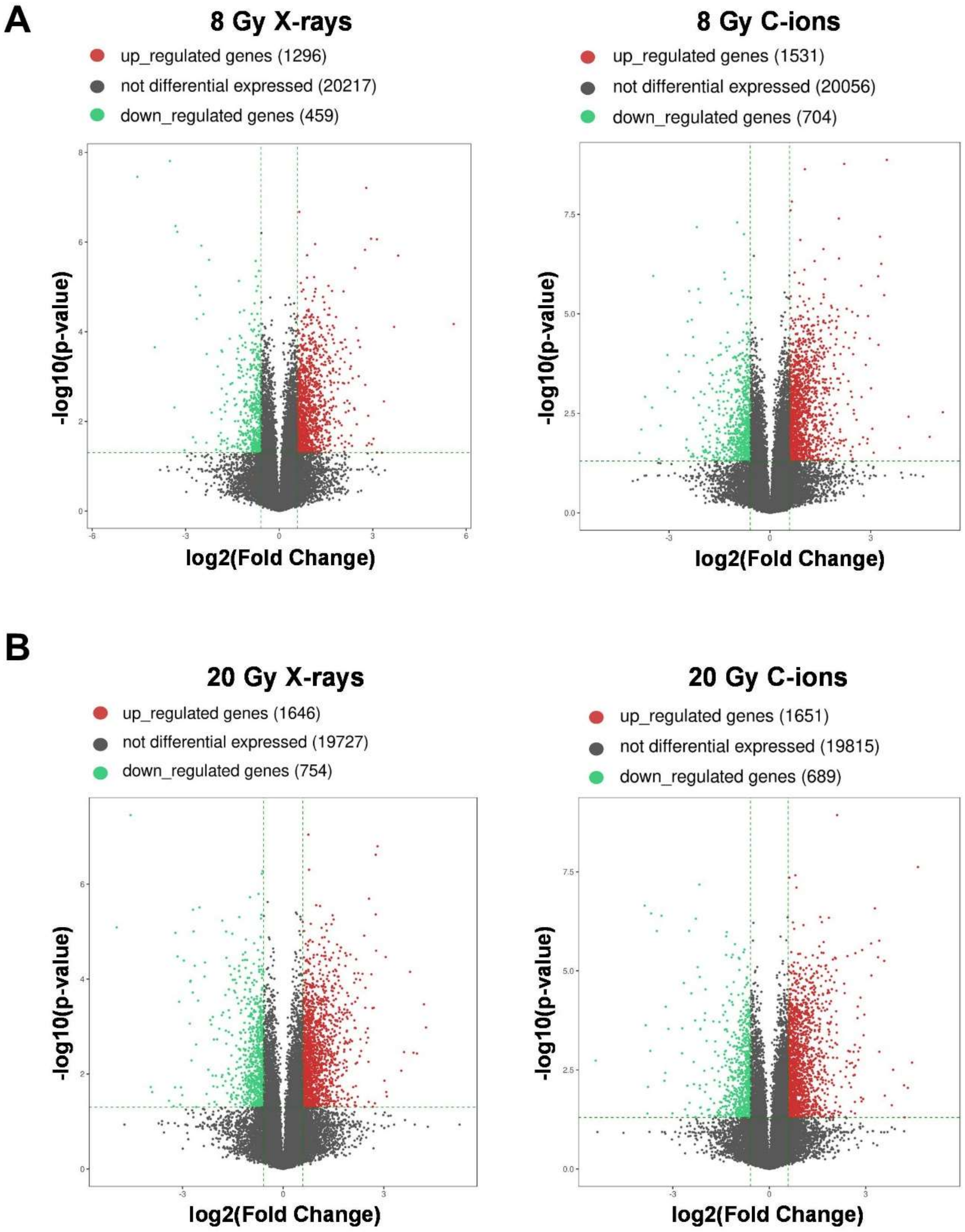

**Supplementary Figure 3**. **Differential expression of genes for various doses of X-rays in 4T1 and TS/A cells**. Volcano plots of differentially expressed genes at 24h after 8 Gy (A) or 20 Gy (B) irradiation with X-rays (left) or carbon ions (right) compared to the respective non-irradiated samples (0 Gy). Significantly upregulated genes and downregulated genes are depicted in red and green, respectively, and non-significant genes are shown in gray. Increasing the dose, the number of differentially expressed genes becomes comparable between the two radiation qualities.

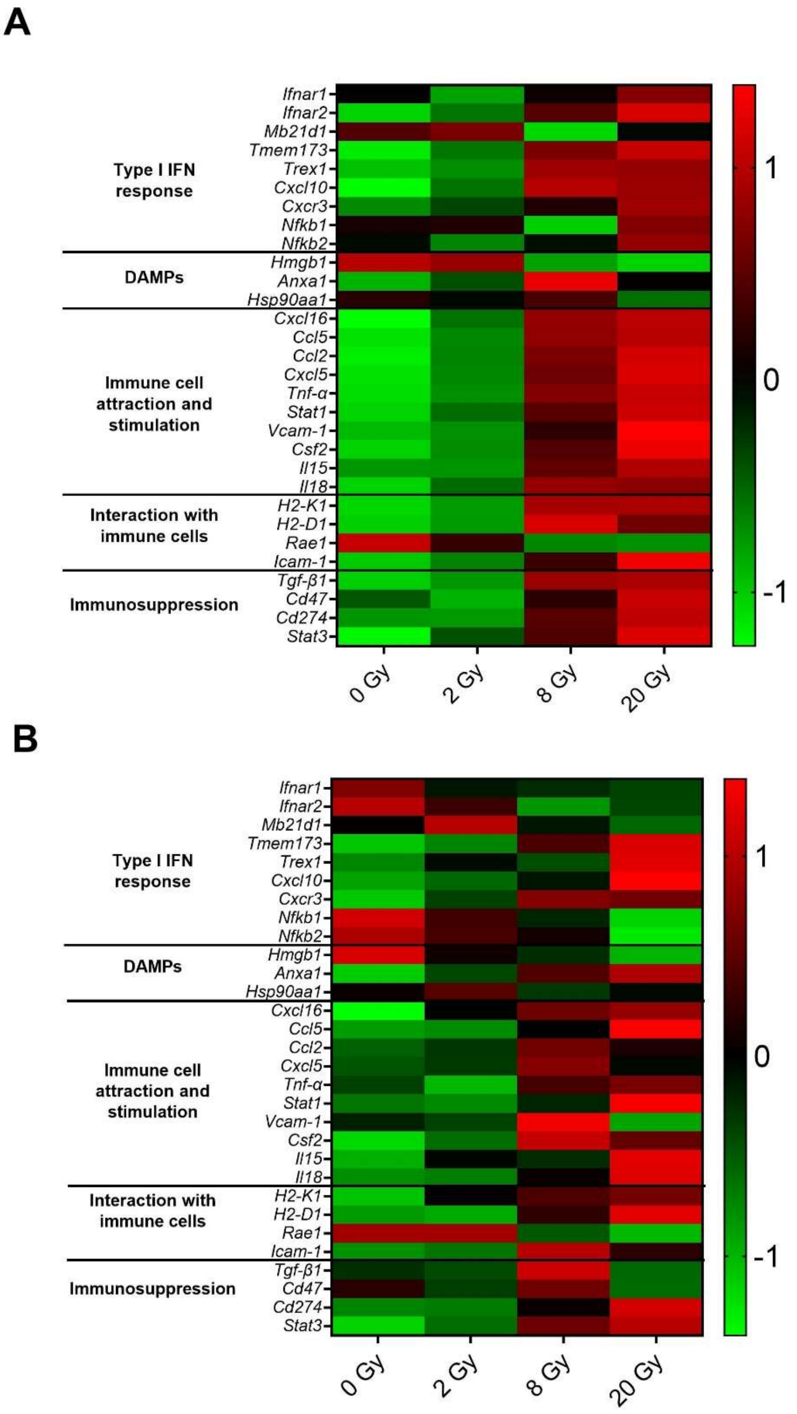

**Supplementary Figure 4**. **Differential expression of selected genes for various doses of X-rays in 4T1 and TS/A cells**. The plot depicts the Z-score hierarchical clustering heat map visualization of the expression of some genes of interest (rows) across different doses (columns) for 4T1 (A) and TS/A (B) cells. Colors represent scaled expression levels, with green for low expression and red for high expression levels.

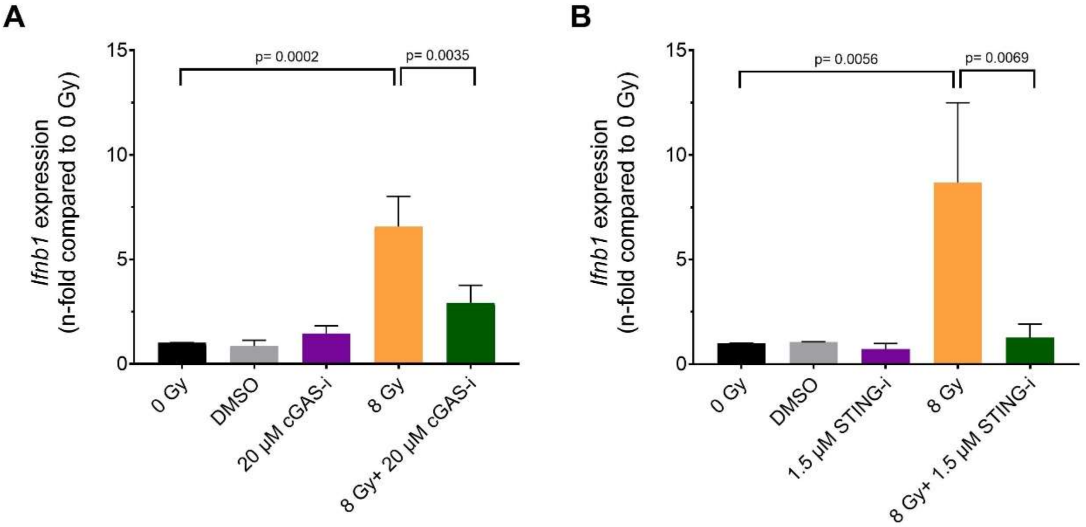

**Supplementary Figure 5**. **Selective inhibition of cGAS or STING diminishes the expression of *Ifnb1* in 4T1 cells.** While exposure to 8 Gy X-rays results in an increased expression of *Ifnb1*, the effects are diminished applying selective inhibitors for cGAS (cGAS-i, A) or STING (STING-i, B) together with radiation exposure.

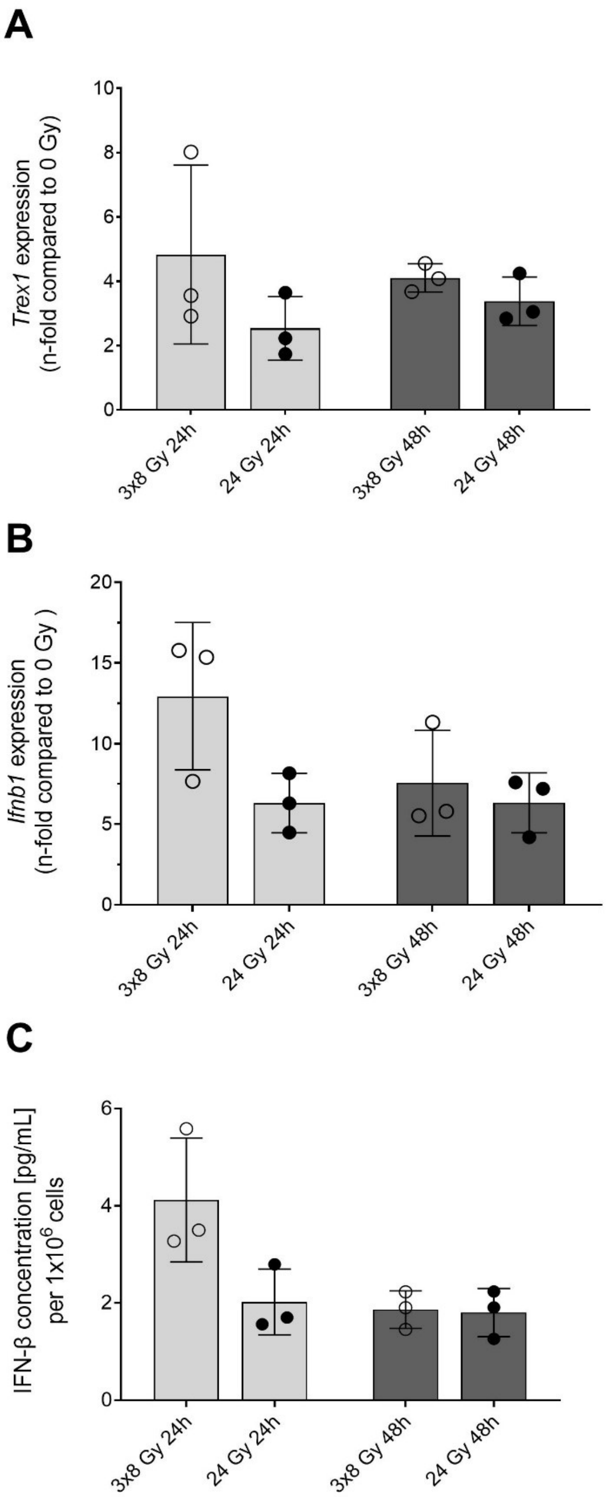

**Supplementary Figure 6. Comparison between single high doses and a 3x8 Gy hypofractionation scheme in TS/A cells.** The endpoints tested for the dose response curves were also performed comparing a single high dose of 24 Gy to a 3x8 Gy hypofractionation scheme at 24h and 48h after exposure to X-rays. More in detail, *Trex1* expression (A), *Ifnb1* expression (B), and IFN-β release (C) were measured. Significances were tested using an unpaired two-tailed t-test. None of the studied endpoints showed significant differences between single dose and fractionated dose.

### **Supplementary Methods:**

Bulk RNA Sequencing

Total RNA from 4T1 and TS/A cells was quantified using a NanoDrop ND-1000 instrument. 1 to 2 µg of total RNA was used to prepare the sequencing library: the total RNA was enriched by oligo (dT) magnetic beads (rRNA removed); RNA-seq library was prepared using KAPA Stranded RNA-Seq Library Prep Kit (Roche), for a strand-specific RNA-seq library. The completed libraries were qualified with Agilent 2100 Bioanalyzer and quantified by absolute quantification qPCR method. To sequence the libraries on the Illumina NovaSeq 6000 instrument, the barcoded libraries were mixed, denatured to single stranded DNA in NaOH, captured on Illumina flow cell, amplified in situ, and subsequently sequenced for 150 cycles for both ends on Illumina NovaSeq 6000 instrument. Image analysis and base calling were performed using Solexa pipeline v1.8 (Off-Line Base Caller software, v1.8). Sequence quality was examined using the FastQC [1] software (version 0.11.7). The trimmed reads (trimmed 5', 3'-adaptor bases using cutadapt [2] version 1.17) were aligned to reference genome using Hisat2 software [3] (version 2.1.0). The transcript abundances for each sample was estimated with StringTie [4] (version 1.3.3), and the FPKM alue for gene and transcript level were calculated with R package Ballgown [5–8] (version 2.10.0). The differentially expressed genes and transcripts were filtered using R package Ballgown. Fold change (cutoff 1.5), p-value (≤ 0.05) and FPKM (≥ 0.5 mean in one group) were used for filtering differentially expressed genes and transcripts. Principle Component Analysis (PCA) and correlation analysis were based on gene expression level, Hierarchical Clustering, Gene Ontology, Pathway analysis, scatter plots, and volcano plots were performed with the differentially expressed genes in R (version 3.5.0), Python (version 2.7) or shell environment for statistical computing and graphics. The Z-score hierarchical clustering heat maps were obtained for each Gene Of Interest (GOI) with the formula:

$$Z - score = \frac{(FPKM\ sample - FPKM\ base\ mean)}{FPKM\ base\ SD}$$

FPKM: fragments per kilobase of transcript per million mapped reads
FPKM sample: mean of FPKM values for the biological triplicates of the GOI
FPKM base mean: mean of FPKM values across all the samples for the GOI
FPKM base SD: standard deviation of FPKM values across all the samples for the GOI

Best fit of the functions

Data in Figure 2A were fitted by the linear function:

$$Y = \alpha D$$

were Y is the number of cytoplasmic dsDNA foci/cell, α [$Gy^{-1}$] is the slope and D the dose in Gy. The α values were 2.53±0.19 [$Gy^{-1}$] for X-rays and 3.68±0.02 [$Gy^{-1}$] for C-ions. The difference is statistically significant (t-test; $p<0.0001$).

We performed the fit of the experimental data in Figure 2 C-D for *Trex1* expression Y vs. dose D with the function:

$$Y = 1 + (P - 1)(1 - e^{-kD})$$

where k ($Gy^{-1}$) is the slope of the curve, P the plateau value of the *Trex1* expression, D the dose in Gy. The best fit values and their uncertainty are reported in Table S1:

**Table S1**: Fitting parameters for the data in Figure 2 C-D.

| **Time after exposure** | **Radiation** | **k±SD** | **P±SD** |
|---|---|---|---|
| **24 h** | X-rays | 0.10±0.07 | 3.1±1.0 |
| | C-ions | 0.12±0.11 | 4.4±1.9 |
| | p-value | 0.643 | 0.0662 |
| **48 h** | X-rays | 0.16±0.10 | 3.5±0.7 |
| | C-ions | 0.34±0.21 | 5.1±0.6 |
| | p-value | 0.0123 | <0.0001 |

An unpaired two-tailed t-test was used to test the difference of P and k parameters between X-rays and carbon ions´ fit. The p-values are shown in Table S1.

The same function was used to fit *Ifnb1* expression (Figure 4 A, C). However, there is no statistically significant difference between X-rays and C-ions. Therefore, we do not compare fitting parameters.

The IFN-β concentration data for X-rays and C-ions (Figure 4 B, D) seem to follow a different trend, with X-rays still increasing at 20 Gy while C-ions are in plateau. Therefore, we do not compare fitting parameters here.
